# Soil-transmitted helminthiasis in children of a rural community part of a school-based deworming program: a cross-sectional study in the Peruvian Amazon

**DOI:** 10.1101/458661

**Authors:** Renato A. Errea, George Vasquez-Rios, María L. Calderon, Diego Siu, Kevin R. Duque, Luciana H. Juarez, Rodrigo Gallegos, Celene Uriol, Claudia R. Rondon, Katia P. Baca, Rosario J. Fabian, Marco Canales, Angelica Terashima, Luis A. Marcos, Frine Samalvides

**Author notes:** Current address: Department of Global Health and Social Medicine, Harvard Medical School. Boston, Massachusetts, United States of America. Corresponding author: (RAE).

## Abstract

**Background:** Children in the Peruvian Amazon basin are at risk of soil-transmitted helminthiases (STH). The aim of this study was to determine the prevalence of STH (*Ascaris lumbricoides, Trichuris trichiura*, hookworm and *Strongyloides stercoralis*) in children from a rural community in the Peruvian Amazon and associated clinical, maternal, sanitation and hygiene factors associated. The community had an active school-based deworming program with mebendazole.

**Methods:** A cross-sectional study was carried out in children aged 2–14 years in Iquitos, Peru; by parasitological analysis of their stools through five methods: direct smear examination, Kato-Katz, spontaneous sedimentation in tube, Baermann method modified by Lumbreras and agar plate culture. Mothers of the participating children were also invited to participate in the study. A questionnaire was completed by each participant to collect demographic and epidemiological information of interest.

**Results:** Among 124 children, 25.8% (32/124) had one or more STH. Prevalence of A. *lumbricoides* was 16.1% (20/124); *S. stercoralis*, 10.5% (13/124); hookworm, 1.6% (2/124) and *T. trichiura*, 1.6% (2/124). STH in mothers was higher in children with any STH than in children without any STH (36.4% vs 14.1%, p<0.02). Prevalence of the common STH (*A. lumbricoides, T. trichiura* and hookworm) was higher in children aged 2–5 than in older children (31.6% vs 12.8%; p=0.01). Several hygiene and sanitation deficits were identified; of which walking barefoot was significantly associated with STH infection (OR= 3.28; 95% CI= 1.11–12.07).

**Conclusions:** STH are highly prevalent in children in this community; *A. lumbricoides* and *S. stercoralis* infections were the most frequent. Further studies should aim to understand the persistent high prevalence of common STH in endemic areas where massive drug administration is practiced, and to determine the appropriateness of controlling STH in mothers and *S. stercoralis* infection. Walking barefoot and other hygiene and sanitation conditions need to be addressed in this community.

**AUTHOR SUMMARY:** Few studies assessing soil-transmitted helminth infections in children and their risk factors have yet been conducted in the Peruvian Amazon. Even fewer reports exist from areas where mass drug administration programs have been initiated. In this study we provide insight to the frequency of soil-transmitted helminths in a setting with an ongoing school-based deworming program.

Besides the most common soil-transmitted helminths (*Ascaris lumbricoides, Trichuris trichiura* and hookworm), we assessed the prevalence of *Strongyloides stercoralis*. Excluding the latter from intestinal helminths studies have often underestimate its frequency and impact in children.

We also surveyed for helminth infection in the mothers of the participating children as infection in caregivers could theoretically be associated with infection in children as they both may share same environmental and behavioral factors associated with STH infections. To our knowledge, this is the first Peruvian study assessing children and mother infection together.

In addition, our results highlight the suboptimal hygiene and sanitation conditions in which people from this rural community live. It likely represents the situation of other rural Amazonian communities in Peru. Public efforts are needed to provide these populations with dignified living conditions and to follow the equity global health agenda.

## INTRODUCTION

Infections by the soil-transmitted helminths (STH) *Ascaris lumbricoides, Trichuris trichiura, Ancylostoma duodenale*/*Necator americanus* (hookworms) and *Strongyloides stercoralis* disproportionately affect children around the world [1]. Due to their transmission associated with poor sanitary conditions and inadequate hygiene practices, higher burden of disease is seen in children from developing countries from sub-Saharan Africa, Southeast Asia and Latin America [1]. Around 267 million pre-school-age children (PSAC) and 568 million school-age children (SAC) worldwide are in risk of STH infection and impaired child growth and cognitive development from A. lumbricoides, T. trichiura and hookworm infections [2], as well as death due to severe *S.stercoralis* infection [3]. Their control is a global health priority.

Water, sanitation and hygiene (WASH) interventions alongside repeated chemotherapy at regular intervals are tenets for the control of STH [4,5]. Despite the progress made, there are still 49% PSAC and 31% SAC in endemic countries not receiving preventive chemotherapy [6]. Similarly, 2.3 billion people do not have adequate sanitation service, 892 million practice open defecation and 884 million lack basic drinking water service [7]. Scale-up of deworming programs in endemic countries and global outreach of WASH programs are necessary.

The Amazon basin in Peru is in special risk of STH due to its tropical climate (favoring geohelminth survival) and the health inequalities present in the region. In Peru, while health interventions have rapidly increased in poor areas of the Andes, the Amazon has experienced less and slower health progress [8]. Moreover, some antihelminth programs already implemented in the region have reported low chemotherapy drug coverage [9]. Research and evaluation of Amazonian communities are needed. Thus, the objective of this study is to determine the prevalence of STH infection in children in a rural community in the Amazon jungle, as well as to identify demographic, medical, maternal, and sanitation and hygiene factors associated with the infection.

## METHODS

### Study design

We conducted a cross-sectional survey study to determine the point prevalence of *A. lumbricoides, T.trichiura*, hookworm and *S. stercoralis* infections in children aged 2–14 years from Padre Cocha, a rural community in Loreto-Peru, using multiple parasitological diagnostic techniques: direct smear examination, Kato-Katz technique, spontaneous sedimentation in tube technique, Baermann method modified by Lumbreras and agar plate culture. We collected information on children’s past medical history and hygiene practices as well as household sanitation conditions. Also, we explored the presence of STH in mothers of the children tested and examined the association between characteristics of the mothers and children’s STH infection status.

### Study population

The community of Padre Cocha belongs to the district of Punchana, province of Maynas, in the region of Loreto, Peru, and it is located by the Nanay River (one of the main tributaries of the Amazon River), 20 minutes by river to the closest urban area. Its population is about 850 inhabitants, distributed in 195 households [10]. In 2012, around 31% were considered poor and 11% lived in extreme poverty, being the main economic activities fishing, farming, craft-making and the increasing touristic activities in the area. The great majority of inhabitants spoke Spanish as their mother language. The local primary care health center in the community was mainly managed by nursing personnel with the sporadic support of a general medical doctor; services were provided 3–4 hours per day during week days [11].

Since 2012, a school-based deworming program initiated in the region of Loreto. The program consisted on the administration of 500mg of mebendazole on a quarterly basis to children aged 3-17, aimed to reduce the burden of *A.lumbricoides, T. trichiura* and hookworm infections (common STH) [9]. During the time of the study, the deworming program had been in place in the community at least two years. Drugs were provided both in the nursing-school to children 3–5 years, as well as in the elementary, middle and high schools (the three located in the same building), for children 6–17 years old.

### Data and stool collection

Once authorization and support from the community leaders were obtained, households with eligible children were identified: children aged 2–14 and residing in Padre Cocha for at least one year. The aim was to identify as many eligible children as possible within the period of December 2^nd^ and 13^th^ of 2015. For that purpose, every block of the community including those geographically hard-to-reach houses were covered. If parents were not present during the encounter, 1–2 more household visits within 24–72 hours were done. If the child was not present, voluntary informed consent from parents was collected and the child was approached later at the local school. Mothers willing to participate also provided informed consent for their own participation. Only children who assented to participate were included. A questionnaire was administered to collect information on socio-demographic characteristics, relevant medical history, hygiene practice and household sanitation of each child, as well as socio-demographic characteristics of mothers. Weight and height in children were measured using calibrated instruments. At the end of the interview, two stool containers were given to participants. They were detailly explained to provide the child and mother’s stool samples with at least a 24-hour difference between samples and to deposit them immediately in the local health center. The research team collected the stool samples every 6–12 hours at the local health center and immediately sent them for analysis at Hospital Regional de Iquitos’ facilities (45 minutes away from the community).

### Parasitological analyses

A senior laboratory technician with over 35 years of experience from Instituto de Medicina Tropical “Alexander Von Humboldt” from Universidad Peruana Cayetano Heredia, performed the parasitological analysis at Hospital Regional de Iquitos’s laboratory. Multiple parasitological techniques were employed to increase diagnosis performance. Each sample was analyzed through direct smear examination, spontaneous sedimentation in tube technique, Kato-Katz, Baermann method modified by Lumbreras (BM) and agar plate culture. Spontaneous sedimentation in tube and Kato-Katz were performed mainly for detection of *A. lumbricoides, T. trichiura* and hookworm infection [12], while Baermann technique and agar plate culture were employed for detecting *S. stercoralis* infection [13]. All the techniques were performed following the guidelines for intestinal parasites diagnosis from the National Institute of Health of Peru [14].

### Sample size calculation and statistical analysis

Of the total population of 850 inhabitants in Padre Cocha [10], approximately 32% are less than 15 years old per local estimates [15]. Considering an estimated frequency of any STH of 50% by December 2015 [9] and a 5% margin of error, the estimated sample size was 136 children, calculated using Epilnfo™.

All data was entered in a Microsoft Excel spreadsheet and posteriorly managed using the statistical package STATA v14 (license: Universidad Peruana Cayetano Heredia). Prevalence comparisons were made using chi-square test or fisher exact test when appropriate, considering a significance level of 0.05. Bivariate analysis with odds ratio calculation was performed for evaluating association with geohelminths infection and the characteristics of interest, considering a 95% confidence interval. Prevalence of STH was considered as the prevalence of any STH. Missing data was reported when >5%.

### Ethics statement

This study was approved by the Institutional Review Boards of Universidad Peruana Cayetano Heredia (#65959) and Hospital Cayetano Heredia (#126-015). Participating children assented to participate in the study and parents provided written informed consent for their children’s participation. Mothers also provided written informed consent for their own participation.

## RESULTS

Between December 2^nd^ and 13^th^, 2015, stool samples from 124 children were collected; 35 children provided two stool samples from separate days, the remainder provided one sample. The mean age was 7. ±3.4 years, 30.7% (38/124) were pre-school-aged children (PSAC, 2–5 years) and 69.3% (86/124) were school-aged children (SAC, 6–14 years); 53.2% (66/124) were males.

Anthropometric measures of children showed that 60.8% (73/120) had stunting according to WHO criteria. Twenty-three percent (40/174) of children reported to have at least three episodes of diarrhea in the last year, 14.6% (18/123) reported to have been regularly absent to school (≥ 4 missed days) in the last year, and 79% (98/123) referred having taken any type of antiparasitic drugs also during the last year. Regarding hygiene practices, 83.9% (104/124) reported irregular handwashing (not washing before meals or after defecation), 66.7% (82/123) walked barefoot, 28.2% (35/124) were nail biters and 20.1% (25/124) used the river for bathing or recreational activities. The 124 children belonged to 78 households, of which 55.6% (40/72) had no running water and 40.8% (31/76) had soil floor; however, 97.4% (75/77) had at least a latrine for defecating.

The total point prevalence of any STH in children was 25.8% (32/124), being A. *lumbricoides* the most frequent (16.1%, 20/124), followed by *S. stercoralis* (10.5%, 13/124); 1.6% (2/124) had hookworm infection and 1.6% (2/124) were infected by *T. trichiura* (Table 1). The prevalence of STH infection was higher in PSAC than in SAC but not statistically significant (36.8% [14/38] vs 20.9% [18/86]; p=0.06); when analyzed only for the common STH (*A. lumbricoides, T. trichiura* and hookworm), the prevalence was significantly higher in PSAC than in SAC (31.6% [12/38] vs 12.8% [11/86]; p=0.01). Regarding the characteristics evaluated for association with STH infection, having taken antiparasitic drugs in the last three months (0R=0.35 [CI= 0.13–0.99]; p=0.02;) showed to be a protector for being infected with STH, while walking barefoot (0R=3.28 [CI= 1.11–12.07]; p=0.02) was a risk factor for STH infection (Table 2).

**Table 1.** Prevalence of soil-transmitted helminths in 124 children in Padre Cocha

|  | No | % |
| --- | --- | --- |
| *Ascaris lumbricoides* | 20 | 16.1 |
| *Strongyloides stercoralis* | 13 | 10.5 |
| *Necator americanus* /<br>*Ancylostoma duodenale* | 2 | 1.6 |
| *Trichuris trichiura* | 2 | 1.6 |
| *Any STH* | 32 | 25.8 |

STH, soil-transmitted helminth

**Table 2.** Association between soil-transmitted helminth infection and medical history, hygiene practices and household sanitation characteristics in 124 children in Padre Cocha.

| Characteristic | STH infection<br>(n=32)<br>No (%) | No STH infection<br>(n=92)<br>No (%) | OR | 95% CI | p |
| --- | --- | --- | --- | --- | --- |
| Use of antiparasitic drugs in the last year | 21/32 (65.6) | 77/91 (84.6) | 0.35 | 0.13-0.99 | 0.02 |
| ≥3 episodes of diarrhea in the last year | 8/32 (25) | 23/91 (25.3) | 0.99 | 0.34-2.68 | 0.98 |
| Chronic malnutrition | 19/31 (61.3) | 54/89 (60.7) | 1.03 | 0.41-2.63 | 0.95 |
| School absenteeism ≥4 days in the last year | 5/31 (16.1) | 13/91 (14.3) | 1.15 | 0.29-3.88 | 0.78 |
| Walking barefoot | 26/31 (83.9) | 56/92 (60.9) | 3.28 | 1.11-12.07 | 0.02 |
| Irregular hand washing | 24/32 (75) | 80/92 (87) | 0.45 | 0.15-1.44 | 0.11 |
| Use of river | 12/31 (38.7) | 31/92 (33.7) | 1.24 | 0.48-3.11 | 0.61 |
| Nail biting | 7/32 (21.9) | 28/92 (30.4) | 0.64 | 0.21-1.76 | 0.35 |
| Soil floor | 15/32 (46.9) | 36/91 (39.6) | 1.35 | 0.55-3.28 | 0.47 |
| Open defecation | 0/32 (0) | 1/90 (1.1) | -- | -- | -- |
| No running water | 16/31 (51.6) | 51/88 (58) | 0.77 | 0.31-1.92 | 0.54 |

STH, soil-transmitted helminth

In addition, we collected stool samples from 51 mothers, who represented the mothers of 69.4% (86/124) of the children tested. Mean age of mothers was 35.4 ± 9.2 years, 25.5% (13/51) were single mothers and 59.2% (29/49) had not completed high school. Prevalence of STHs in mothers was 19.6% (10/51); 9.8% (5/51) were infected by *A. lumbricoides*, 5.9% (3/51) by *S. stercoralis*, 3.9% (2/51) by hookworms and none by *T. trichiura*. There was not significant statistical difference between single marital status, low educational level (less than high school) or young age (≤25 years) of the mother and STH infection in children (Table 3); however, prevalence of STH infection in mothers was significantly higher in children infected than in the non-infected (36.4% [8/22] vs 14.1% [9/64], p<0.02).

**Table 3.**
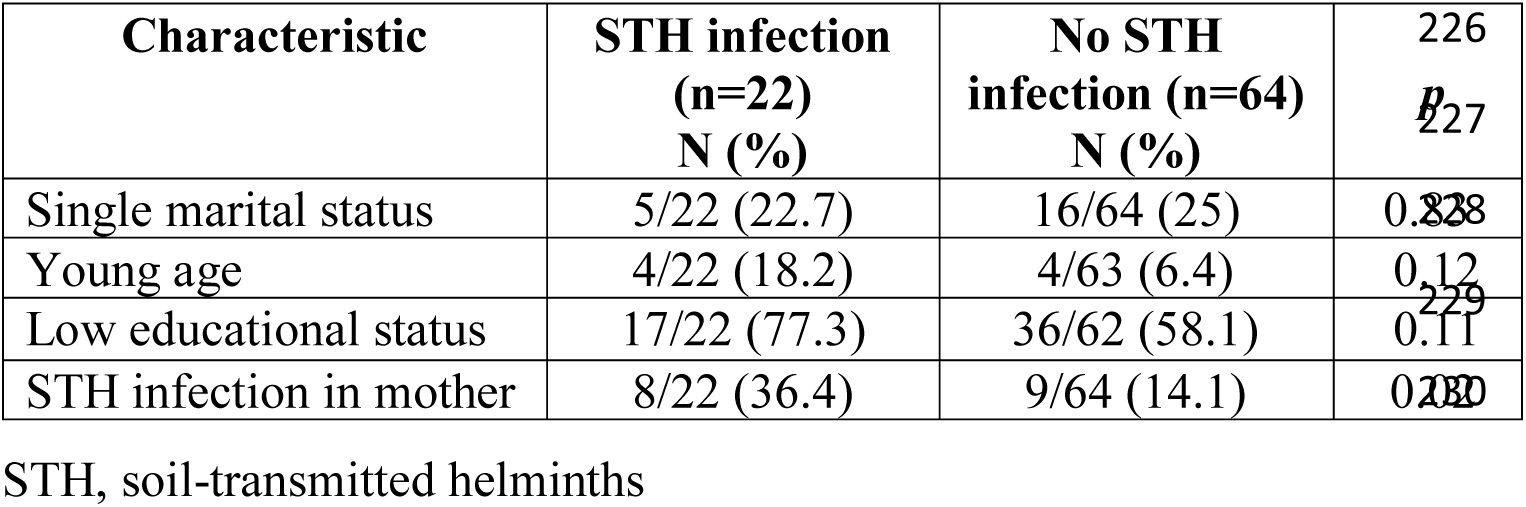
Association between characteristics of the mothers and soil-transmitted helminth infection in children (n=86) in Padre Cocha

## DISCUSSION

A 26% prevalence of STH was found in children from an Amazonian rural community part of a school-based deworming program. Most of the burden came from *A. lumbricoides* infection (16.5) - high despite the delivery of mebendazole every three months - and from *S. stercoralis* (10.5%) - similar prevalence compared to another local study [16]. In the other part, *T. trichiura* and hookworm infections were not frequent (1.6%) and lower compared to other similar studies conducted in the Peruvian jungle [9,12,17,18].

The prevalence of the common STH (*A. lumbricoides, T. trichiura* and hookworm) was higher in PSAC (31.6%) than in SAC (12.8%). This is of special importance because recent evidence has shown robust positive effects of deworming in PSAC [19]. Thus, further studies should evaluate the reasons behind these dissimilar prevalence rates. In that regards, it is necessary to take into consideration that, in Padre Cocha, the nursing school (for PSAC) was located and managed separately from the upper-level schools (elementary, middle and high school were led by the same director in the same territory), requiring separate coordination and surveillance procedures. This may have resulted in different practices, supply and/or drug coverage, as has been reported in the literature [20].

*S. stercoralis* infection was a frequent infection in children of this community. Strongyloidiasis has been frequently associated with diarrhea and abdominal pain in the pediatric population [21], being even associated with stunting in children with severe infection [22]. Nonetheless, mebendazole, the drug administered as part of the deworming program, is not effective against *S. stercoralis* [23]. Thiabendazole and ivermectin have shown to be effective within strongyloidiasis control programs [24-26]. Thus, research evaluating the appropriateness of adding these drugs in Padre Cocha and alike communities in Peru is needed.

In this study the prevalence of STH infection in mothers was higher in children infected (36.4%) than in the non-infected (14.1%). Mother helminth infection has been reported as a risk factor for infection in children [27]; therefore, it could be contributing to the high prevalence of STH in children of this community. Currently, the WHO recommends administering antiparasitic drugs not only to children but also to women of childbearing age [5]. Following these study’s results and WHO recommendations, providing anti-helminths for mothers should be considered.

A great percentage of the population lacked optimal sanitary conditions and adequate hygiene practices in Padre Cocha. Walking barefoot was the only found risk factor for infection by STH, yet 87% of the population had irregular hand washing habits, 55% percent of the households did not have running water and 39.7% had soil floor; all of them reported in the literature as risk factors [4,28]. To combat this, a hygiene education intervention in another community in the Peruvian jungle was effective in reducing children helminth infections [18]. Padre Cocha and other rural Amazonian communities urge similar interventions and sustained public efforts for improving access to clean water, hygiene and sanitation conditions.

Over half of the pediatric population studied had stunting, proportion substantially higher than the 17.3% National estimate [8]. Although stunting was not associated with STH infection in this study, several studies have reported the effect of moderate-to-high infection rates on child nutritional status [29]. Moreover, low nutritional status also influences the impact of the infection [29]. As has also been found by neighbor countries [30], rural Amazon communities may be at special risk of children stunting, which requires public health attention.

Finally, taking antiparasitic drugs in the last three months was associated with reduced infection by any STH. However, the way the information was collected could not specify if the drug taken was the one provided by the school-based deworming program. In fact, the health center was stacked, although limitedly, with anthelmintic drugs and other parasitic medicine. Further studies on the drugs available for this community could help elucidate this association.

This study had several limitations. First, it was not possible to collect more than one stool sample from all the participants; however, despite the several challenges that conducting on-field studies in resource-constrained settings involve, recommended, multiple parasitological techniques for these scenarios were employed, improving the detection capacity of parasites [12,13]. Second, not all households could be reached despite coming back up to two more times to each house where nobody was initially present. Third, samples from all mothers of participating children could not be obtained, perhaps because mothers were more motivated to find if their children were infected rather than themselves; nonetheless, participating mothers were the mothers of almost 70% of the participating children. Finally, we could not conduct multivariate analysis and adjust for confounding due to not enough sample size; however, results could be generalized to other similar rural communities of the region.

In conclusion, STH are highly prevalent in children in Padre Cocha with *A. lumbricoides* and *S. stercoralis* infections being the most frequent. Further studies need to be conducted to understand the higher prevalence of common STH in PSAC, as well as to determine the suitability of including treatment for STH in mothers and for *S. stercoralis* infection as part of the regional deworming program. Padre Cocha and alike rural populations deserve a comprehensive program for helminthiases control that not only includes the delivery of antihelminth drugs but the improvement of household conditions and hygiene practices.

## ACKNOWLEDGEMENTS

We are grateful to the participants for their kind, voluntary participation and to the community authorities for their support to the study. We also want to thank Carmen Quijano and Matilde Quijano from the Laboratory of Parasitology of the Instituto de Medicina Tropical “Alexander von Humboldt” for their support with specimen analysis.

